# *Vegfr3* is required in the *Tbx1* expression domain for cardiac outflow tract development

**DOI:** 10.64898/2026.05.29.728669

**Authors:** S. Martucciello, M. Bilio, S. Cioffi, M. Cavallaro, A. Baldini, E. A. Illingworth

**Author notes:** **Corresponding authors** Stefania Martucciello, Ph.D, Associate Professor of Molecular Biology, Department of Chemistry and Biology, University of Salerno, Via Giovanni Paolo II, 132, 84084 Fisciano (SA), Italy, Tel: +39 089/968125, Elizabeth Illingworth, Ph.D, Professor of Molecular Biology, Department of Chemistry and Biology, University of Salerno, Via Giovanni Paolo II, 132, 84084 Fisciano (SA), Italy, Tel: +39 089/968121. equal contribution (M. Bilio, S. Cioffi).

## Abstract

Gene inactivation in model organisms has identified numerous genes and signaling pathways involved in mammalian cardiac OFT development. Human genetics data have implicated the *VEGFR3* gene in OFT development but when and where it is required is unknown. In this study we determined the sensitivity of the developing murine cardiac OFT to reduced *Vegfr3* gene dosage and we tested whether its’ requirement is dependent upon TBX1, a known regulator of *Vegfr3* expression in cardiac and lymphatic endothelial cells. We found that in the mouse, a single copy if the *Vegfr3* gene was sufficient for normal cardiac OFT development in most cases. Mutation of a single copy of the *Tbx1* gene greatly enhanced the sensitivity of OFT development to *Vegfr3* dosage reduction and led to the formation of severe OFT anomalies. In addition, deletion of *Vegfr3* in the *Tbx1* expression domain also led to OFT abnormalities. We used RNAscope to reveal the location of *Vegfr3* and *Tbx1* transcripts in midterm mouse embryos. This revealed co-localization of these transcripts that was restricted to the aortic sac endothelium, suggesting that the distal OFT is a potential site of genetic interaction between *Vegfr3* and TBX1 that is critical for normal OFT development.

## INTRODUCTION

Mutations of the *VEGFR3* and *TBX1* genes are associated with Milroy syndrome and DiGeorge syndrome, respectively. Both diseases affect the vascular system. Milroy syndrome is the most frequent primary hereditary lymphedema. Studies in animal models have shown that *VEGFR3* is a major lymphangiogenesis gene, being required for the formation of the jugular lymph sacs and systemic lymphatic vessels (Karkkainen et al., 2000; Mäkinen et al., 2001). Prior to the development of the lymphatic system, *Vegfr3* is required in blood vessels and its’ loss causes early embryonic lethality due to the failure of remodeling of the primitive vascular plexus, which requires angiogenic sprouting (Dumont et al., 1998). DiGeorge syndrome (DGS) on the other hand, which is primarily a *TBX1* haploinsufficiency disorder, is not commonly associated with lymphatic anomalies, although rare cases have been reported (Mansir et al., 1999; Unolt et al., 2018). In humans, DGS is mainly caused by a heterozygous chromosomal microdeletion on chromosome 22 that encompasses over 40 genes, including *TBX1*. Heterozygous *TBX1* point mutations recapitulate most of the physical defects associated with the 22q11.2 chromosomal microdeletion, including congenital heart defects, skeletal defects, facial dysmorphism and immune defects, thus providing evidence of the pleotropic functions of TBX1 during embryonic and fetal development.

Studies in the mouse have shown that TBX1 plays a critical role in cardio-pharyngeal mesoderm (CPM), a population of progenitors that differentiates into different cell types that contribute to the formation of the heart, facies and neck glands, thereby accounting for the aforementioned clinical signs and symptoms of DGS. Furthermore, homozygous inactivation of *Tbx1* causes generalized lymphatic vessel hypoplasia or aplasia, which is lethal perinatally (Chen et al., 2010); TBX1 activates *Vegfr3* expression in murine lymphatic endothelial cells by binding a putative enhancer in the endogenous *Vegfr3* gene (Chen et al., 2010), and the two molecules interact strongly during cardiac lymphangiogenesis (Martucciello et al., 2020). TBX1 also regulates *Vegfr3* in venous endothelial cells and brain microvessels during murine embryogenesis (Cioffi et al., 2014, 2022).

In humans, rare variants of the *VEGFR3* gene are associated with cardiac outflow tract (OFT) abnormalities, including Tetralogy of Fallot (Jin et al., 2017; Reuter et al., 2019; Page et al., 2019), suggesting a previously unknown role for the gene in cardiac development. Interestingly, Tetralogy of Fallot is the most common congenital heart defect found in patients with DGS. We hypothesize that interaction between TBX1 and *Vegfr3* is required in cardiac progenitors in the CPM, or in their descendents, for development of the cardiac outflow tract (OFT). Studies conducted in mice, including mouse models of DGS, show that TBX1 is required for the deployment of cardiac progenitors from the second heart field (part of the CPM) into the developing OFT (Xu et al., 2004; Rana et al., 2014). This progenitor population gives rise to cardiomyocytes, endothelial cells and branchiomeric myocytes (Milgrom-Hoffman et al., 2011; Devine et al., 2014; Lescroart et al., 2014). Fate mapping experiments have revealed that endothelial cells in the aortic sac (AS) and OFT are of second heart field (SHF) origin (Cai et al., 2003; Verzi et al., 2005) but they have also revealed the existence of different populations of SHF progenitors that are spatially and temporally separate and give rise to different structures and cell types. For example, endothelial cells of the anterior pharyngeal arch arteries (PAAs) and aortic sac arise earlier than those of the posterior PAAs and from a different population of SHF progenitors (Devine et al., 2014; Wang et al., 2017). Thus, while SHF-derived endothelial cells populate the OFT, to our knowledge, it is not known whether they play a critical role in cardiac morphogenesis, or whether Tbx1-fated endothelial cells populate the portion of the OFT involved in septation. We investigated this here using mouse genetics approaches.

## RESULTS

### *Vegfr3* heterozygosity is not a significant cause of OFT maldevelopment

To understand whether *Vegfr3* has a role in murine cardiac development, we first evaluated the effect of heterozygous inactivation in the mouse. We analyzed the cardiac phenotype in coronal sections of isolated embryonic hearts at the fetal stage (E)18.5. The results revealed a low penetrance (10%, 1/11) of peri-membranous intraventricular septal defects (VSD) in *Vegfr3*^*+/-*^ embryos compared to WT (0/5) (Table 1, Figure 1). No other intracardiac or great vessel defects were identified and the cardiac valves appeared to be normal. We have reported mild cardiac lymphatic vessel hypertrophy in *Vegfr3*^*+/-*^ embryos (Martucciello et al., 2020), indicating that in the mouse, loss of one copy *Vegfr3* affects cardiac lymphatic development but not cardiac morphogenesis.

**Table 1.** Cardiac phenotype in *Tbx1*^*Cre/+*^ *;Vegfr3*^*+/-*^ germline embryos (E18.5).

| Genotype (E 18.5) | n | Normal | Cardiac Phenotype |
| --- | --- | --- | --- |
| *WT* | 5 | 5 | None |
| *Tbx1<sup>Cre/+</sup>* | 8 | 8 | None |
| *Tbx1<sup>Cre/+</sup> ; Vegfr3<sup>+/-</sup>* | 8 | 1 | 7 (VSD) |
| *Vegfr3<sup>+/-</sup>* | 11 | 10 | 1 (VSD) |

**Figure 1.**
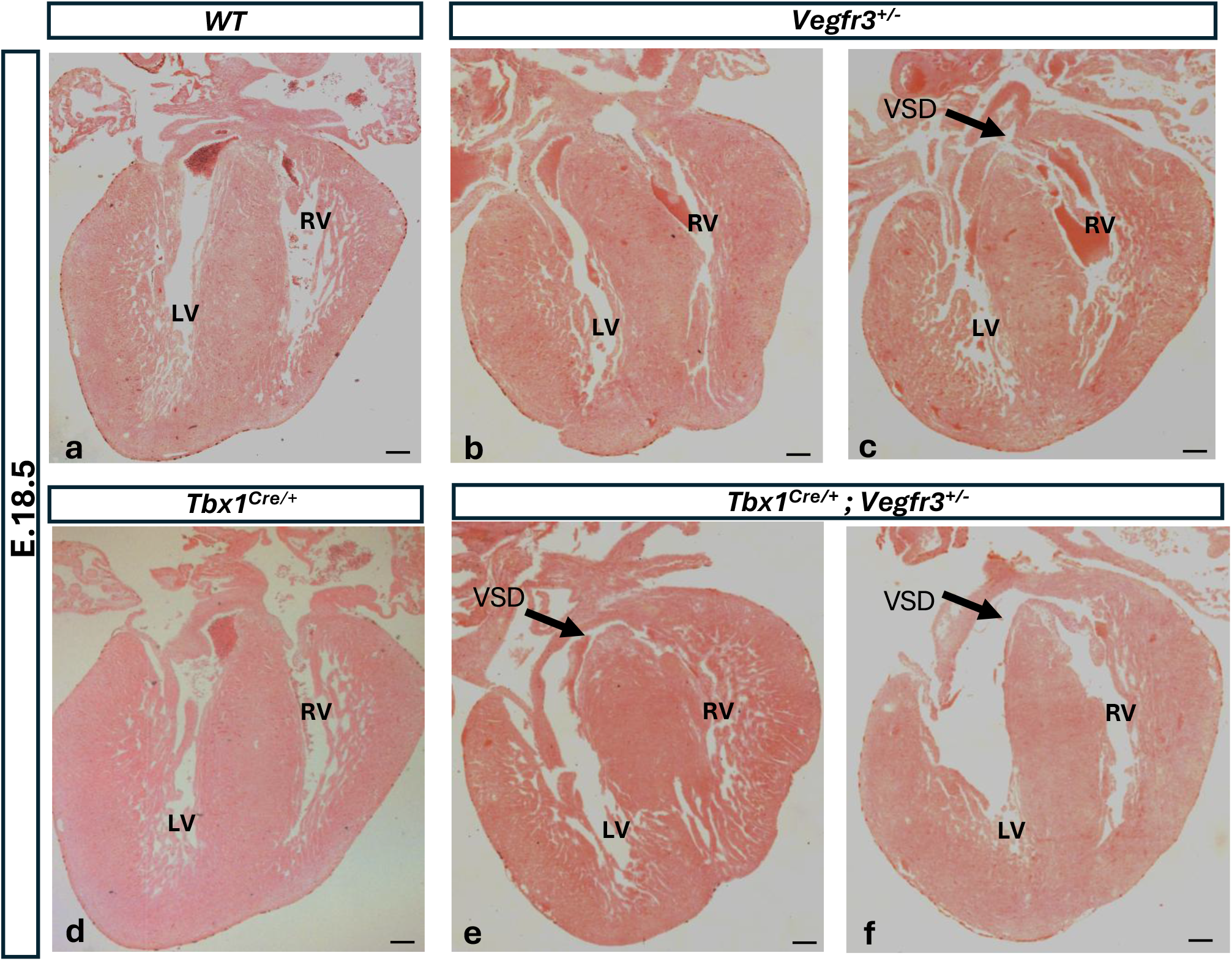
Tbx1 and *Vegfr3* interact genetically in cardiac OFT development. Coronal sections of isolated hearts of *WT* (a), *Vegfr3*^*+/-*^ (b-c), *Tbx1*^*Cre/+*^ (d) and *Tbx1*^*Cre/+;*^; *Vegfr3*^+/-^ (e-f) embryos (E18.5) stained with eosin. Black arrows in panels c, e and f indicate the position of a ventricular septal defect. RV, right ventricle; LV, left ventricle; VSD, ventricular septal defect. Scale bar 100 µm.

### TBX1 and *Vegfr3* interact genetically in cardiac OFT development

We have previously shown that in the mouse TBX1 and *Vegfr3* interact strongly in brain vascularization and during systemic and cardiac lymphangiogenesis (Chen et al., 2010; Cioffi et al., 2014, 2022; Martucciello et al., 2020). To test whether they interact in cardiac OFT development we intercrossed *Tbx1*^*Cre/+*^ and *Vegfr3*^*+/-*^ mice and analyzed the cardiac phenotype in coronal sections of isolated embryonic hearts at E18.5 (*Tbx1*^*Cre*^ is a null allele, (Huynh et al., 2007). We found that seven out of eight compound heterozygous embryos analyzed had a peri-membranous VSD compared to only one out of eleven *Vegfr3*^*+/-*^ embryos and none out of eight *Tbx1*^*Cre/+*^ embryos analyzed (Table 1, Figure 1). Thus, *Tbx1* mutation greatly enhanced the penetrance of VSD in *Vegfr3*^*+/-*^ mutants, suggesting the existence of a genetic interaction.

### *Vegfr3* and *Tbx1* are co-expressed in the aortic sac

In order to gain insights into where and when a genetic interaction might occur, we analyzed expression of *Tbx1* and *Vegfr3* by RNA *in situ* hybridization using the RNAscope™ technology (Bio-techne) on whole wild type embryos at E8.5 and E9.5. We then prepared histological sections of the double-stained embryos in order to obtain a detailed view of the distribution of stained cells. Representative embryos are shown in Figure 2. At E8.5, *Vegfr3* was expressed throughout the developing vascular plexus as expected (Watanabe et al., 2019), in fully formed blood vessels, including the PAAs (Figure 2a-b) aortic sac (AS) and outflow tract (Figure 2d-d’’). At this developmental stage, *Tbx1* was expressed bilaterally in the head mesenchyme (Figure 2c, 2d), in the core of the pharyngeal arches (PA) I and II (Figure 2c, 2c’’’) and as previously reported in the pharyngeal endoderm and in the second heart field (Calmont et al., 2009). No double stained cells (yellow) were detected in any part of the embryo. At E9.5, in the rostral half of the embryo, expression of *Vegfr3* and *Tbx1* were detected in all of the aforementioned tissues. In addition, *Tbx1* was expressed in endothelial cells lining the AS, but not in the OFT, in embryos with >27 somites (Figure 2e-2g). These results indicate that there is no expression overlap between *Tbx1* and *Vegfr3* in the rostral half of the embryo prior to E9.5, but this occurs at the level of the AS from E9.5 onwards.

**Figure 2.**
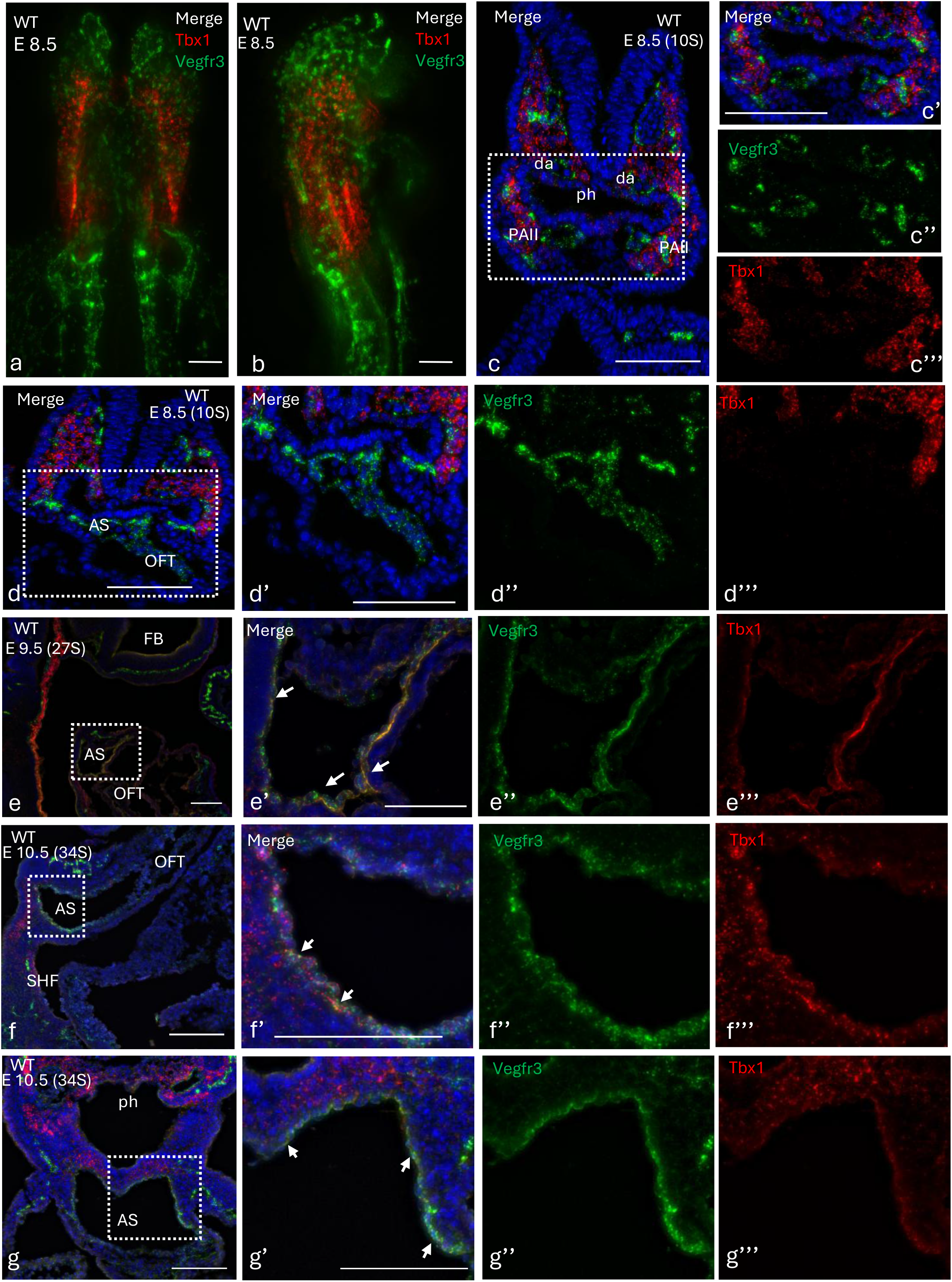
*Tbx1-Vegfr3* co-expression in the aortic sac: (a-b) *Tbx1* (red) and *Vegfr3* (green) expression revealed by RNAscope in wholemount *WT* embryos at E8.5 with a rostro-caudal orientation (ventral (a) and dorsal (b) views), and in transverse sections (c-c’’’) revealed *Vegfr3* but not *Tbx1* expression in the aortic sac. Sagittal sections of *WT* embryos at E9.5 (27 somites) (e-e’’’), E10.5 (34 somites, f-f’’’) and transverse section (34 somites, g-g’’’) showed that *Tbx1* and *Vegfr3* are co-expression in the aortic sac. PAA II, pharyngeal arch artery II; da, dorsal aorta; ph, pharynx; AS, aortic sac; OFT, outflow tract. Scale bar 100 µm.

### *Vegfr3* is required in *Tbx1* expressing cells for normal cardiac outflow tract development

We next tested whether conditional mutation of *Vegfr3* in the *Tbx1* expression domain alone was sufficient to disrupt OFT development. For this, we intercrossed *Tbx1*^*Cre/+*^*;Vegfr3*^*flox/+*^ and *Vegfr3*^*flox/flox*^ mice and analyzed the cardiac phenotype in E18.5 embryos (Table 2, Figure 3). Results showed that *Tbx1*^*Cre*^-driven heterozygous inactivation of *Vegfr3* (thus double heterozygosity in *Tbx1* expressing cells and their descendants) had no effect on OFT development (Figure 3b-b’’). In contrast, the hearts of *Tbx1*^*Cre*^-driven conditional *Vegfr3* homozygous embryos (*Tbx1*^*cre*/+^; *Vegfr3*^*flox*/*flox*^) all presented with one or more cardiac defects (Figure 3c-e), including ventricular septal defects (VSD), atrial septal defects (ASD), double outlet right ventricle (DORV) and overriding aorta (OA). This result indicates that *Vegfr3* expression in the *Tbx1* expression domain is necessary for normal OFT development, and that a single active copy of the *Vegfr3* gene in *Tbx1*-expressing cells, or in their progeny, is sufficient for normal development.

**Table 2.** Cardiac phenotypes in E18.5 embryos with *Tbx1*^*Cre*^-driven conditional inactivation of *Vegfr3*.

| Genotype<br>(E 18.5) | n | Normal | ASD | VSD | DORV | OA |
| --- | --- | --- | --- | --- | --- | --- |
| *Tbx1*<sup>Cre/+</sup> | 8 | 8 | None | None | none | none |
| *Tbx1*<sup>Cre/+</sup> ; *Vegfr3*<sup>flox/+</sup> | 7 | 7 | None | None | none | none |
| *Tbx1*<sup>Cre/+</sup> ; *Vegfr3*<sup>flox/flox</sup> | 7 | None | 4 | 5 | 3 | 2 |

**Figure 3.**
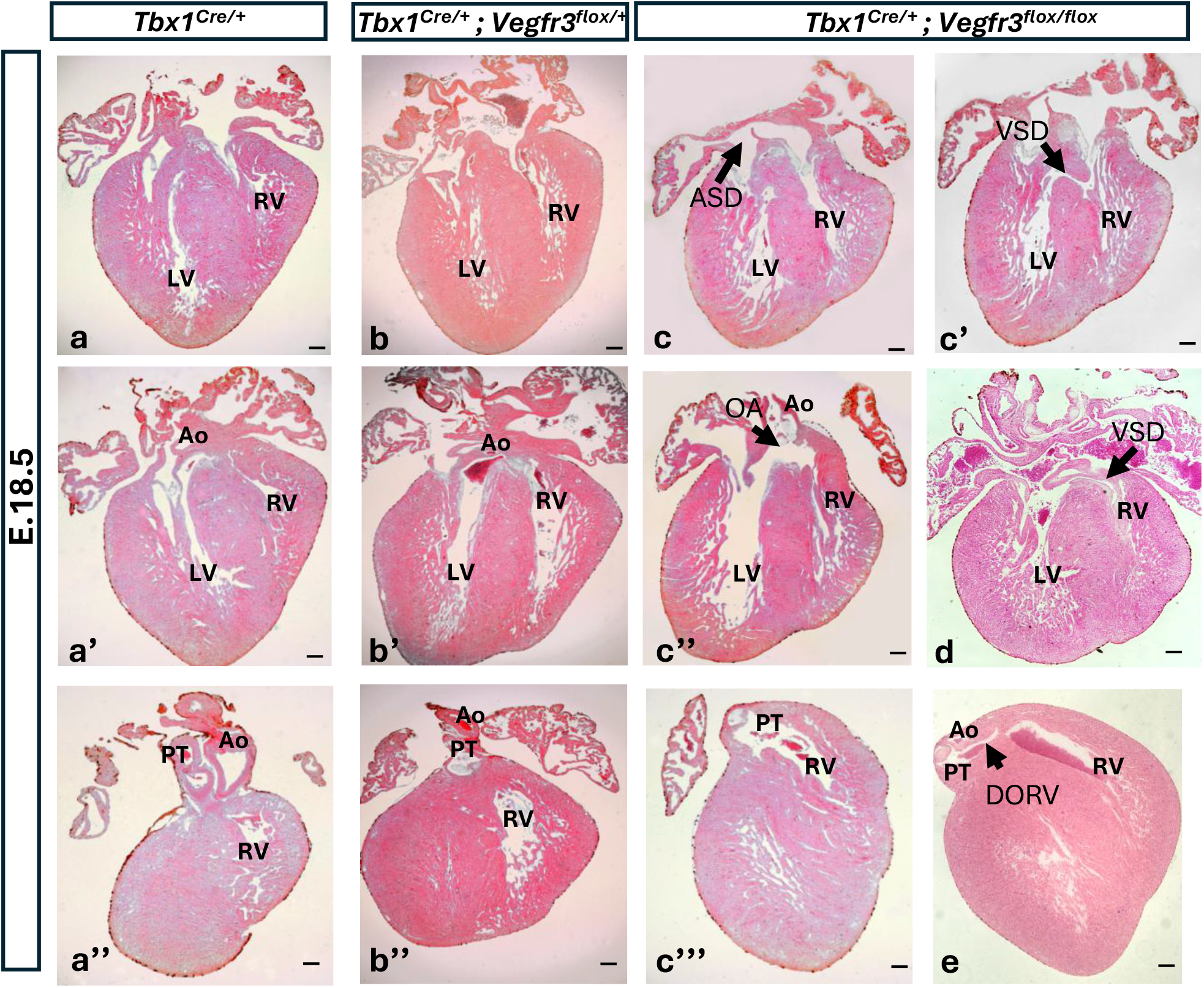
*Tbx1* conditional *Vegfr3* homozygous *embryos* (*Tbx1*^*Cre/+*^; *Vegfr3*^*flox/flox*^*) have* cardiovascular defects. Coronal sections of isolated hearts of *Tbx1*^*Cre/+*^ (a-a’’), *Tbx1*^*Cre/+*;^; *Vegfr3*^*flox/+*^ (b-b’’) and *Tbx1*^*Cre/+*;^; *Vegfr3*^*flox/flox*^ (c-c’’’, d, e) embryos (E18.5) stained with eosin. Black arrows in panels c-c’’, d and e indicate the position of cardiovascular defects. ASD (atrial septal defect); VSD (ventricular septal defect); DORV (double outlet right ventricle); OA (overriding aorta; RV, right ventricle; LV, left ventricle; Ao, aorta; PT, pulmonary trunk. Sections indicated with the same letter are from the same heart. Scale bar 100 µm.

As the endothelium is the only cell type that expresses both *Tbx1* and *Vegfr3*, one would expect that *Tbx1*^*Cre*^-induced recombination of the *Vegfr3*^*flox*^ allele would recapitulate the phenotype observed in compound heterozygotes (Table 1). However, this was not the case, in fact, *Tbx1*^*Cre/+*^*;Vegfr3*^*flox/+*^ mutants survived to adulthood, were fertile and had normal hearts (Table 2). In order to visualize better the extent of *Cre* recombination in endothelial cells we performed double immunofluorescence on sagittal sections of E9.5 *Tbx1*^*Cre/+;*^ *Rosa*^*mTmG*^ embryos using anti-GFP and anti-VEGFR3 antibodies. Results showed that there were many double stained endothelial cells (yellow) in the AS (Figure 4a). Thus, there is overlap between Cre recombination and VEGFR3 expression but evidently, removal of a single copy of the *Vegfr3* gene from this region of overlap is not sufficient to generate OFT defects, even in the presence of *Tbx1* heterozygosity, while removal of both copies is detrimental. We speculate that this may occur because the region of overlap between Cre recombination and *Vegfr3* expression does not fully cover the *Vegfr3* critical domain necessary for normal OFT development. Alternatively, inefficient Cre-driven recombination may not reduce sufficiently the RNA dosage from the heterozygous floxed allele.

**Figure 4.**
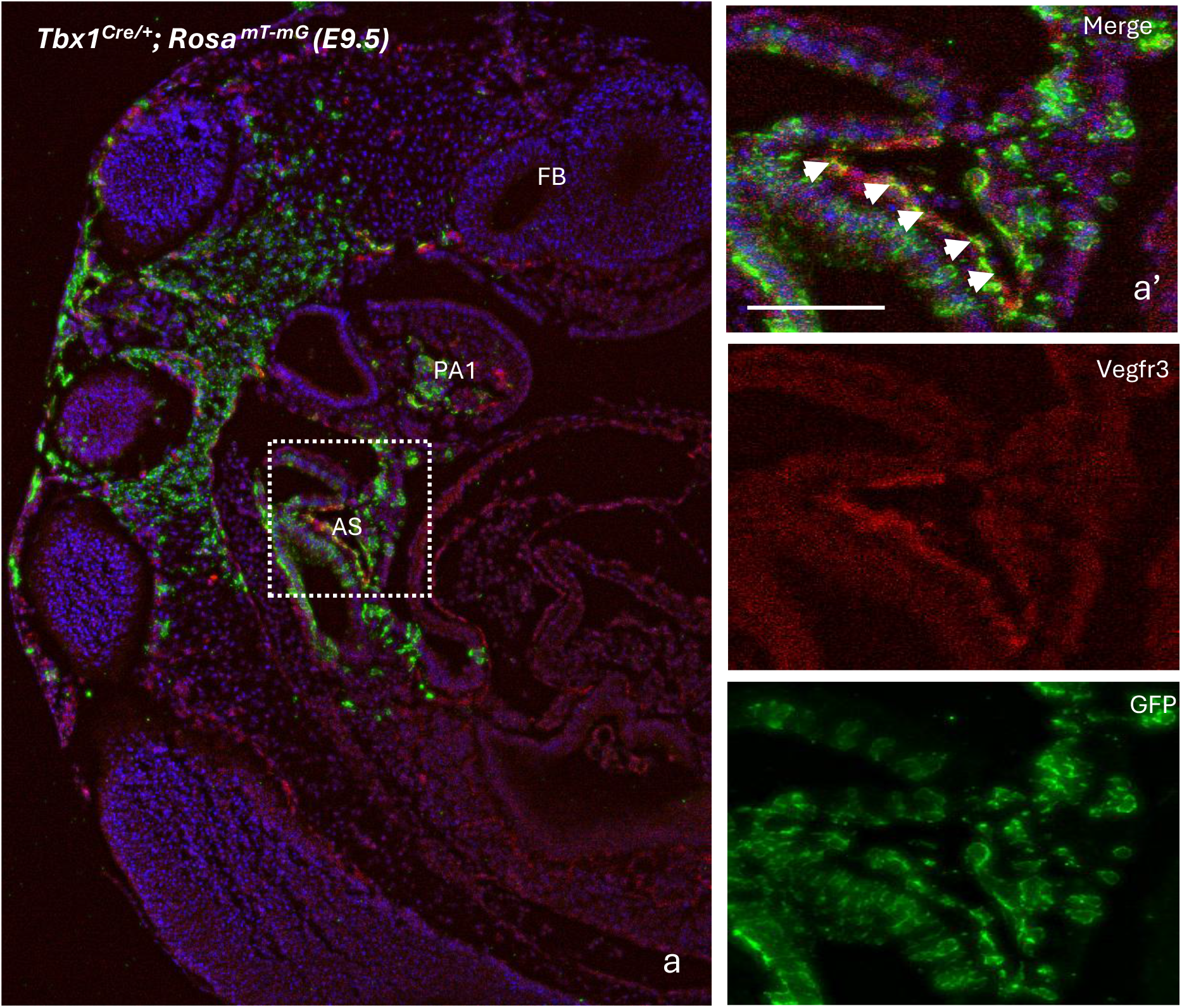
Tbx1 activation in *Vegfr3*-expressing endothelial cells: Anti-Vegfr3 (red) and anti-Tbx1-GFP (green) immunostaining of sagittal sections (a-a’’’) of *Tbx1*^*Cre/+;*^ *Rosa*^*mT-mG*^ reveal Tbx1-activation in *Vegfr3*-expressing endothelial cells of the aortic sac (white arrows). Scale bar 100 µm.

## DISCUSSION

The results of our work indicate that in the mouse, a single copy of the *Vegfr3* gene is sufficient to support normal heart development in most cases, as only a single case (1/11) of VSD was observed in *Vegfr3*^*+/-*^ pre-term embryos. A larger study would be necessary to determine whether 10% is the true frequency of CHDs in *Vegfr3* heterozygotes. In a recent review by Monaghan and colleagues, the authors proposed that TOF in individuals with *VEGFR3* variants or mutations, is not due to *VEGFR3* haploinsufficiency but rather to the presence of truncated VEGFR3 protein during fetal development (Monaghan et al., 2024). This deduction was based on published data, showing that, i) mice carrying *VEGFR3* heterozygous mutations or hypomorphic alleles have normal cardiovascular development (Haiko et al., 2008) ii) in humans, [homozygous?] truncating mutations are very rare, and thus presumed to be mostly lethal (Gordon et al., 2013; Monaghan et al., 2021), iii) in TOF cases with truncating mutations, the mutant protein retained the N-terminal domain but had a shorter C-terminal domain (Reuter et al., 2019). The authors speculated that truncated VEGFR3 proteins disrupt molecular interactions and cellular functions required for normal cardiogenesis (Monaghan et al., 2024). In our study, the *Vegfr3* mutation is inactivating, therefore the presence of truncated proteins can be excluded. Thus, the CHDs observed in *Vegfr3*^*+/-*^ mice here are attributable to gene haploinsufficiency or loss of function. Noteworthy is the observation of *Vegfr3* haploinsufficiency in cardiac lymphatic development (Martucciello et al., 2020), suggesting that haploinsufficiency is context-dependent, where different endothelial populations and subtypes have different sensitivities to *Vegfr3* gene dosage.

Our study showed that heterozygous loss of *Tbx1* greatly enhanced the penetrance of VSDs in *Vegfr3*^*+/-*^ mutants, from 10% to >85% (Table 1). This might be because TBX1 positively regulates *Vegfr3* in some endothelial cells (Chen et al., 2010; Martucciello et al., 2020; Cioffi et al., 2022), therefore its’ mutation might reduce *Vegfr3* expression below a critical threshold required for normal cardiac OFT development, or it might do so indirectly, by disrupting the expression of other *Vegfr3* regulators or by affecting VEGFR3 functions downstream.

Given the strong effect of the *Tbx1* mutation in compound germline *Vegfr3* heterozygotes, we were surprised to find that conditional heterozygous inactivation of *Vegfr3* in the *Tbx1* lineage had no effect on cardiac morphogenesis. We expected that *Tbx1*^*Cre/+*^*;Vegfr3*^*flox/+*^ and *Tbx1*^*Cre/+*^*;Vegfr3*^*+/-*^ embryos would have the same, or similar phenotype, because the *Tbx1*^*Cre*^ allele, which is inactivating (Huynh et al., 2007), is present in both compound genotypes. Therefore, any reduction of *Vegfr3* expression due to the *Tbx1* mutation or to *Tbx1*^*Cre*^-activated recombination of the floxed *Vegfr3* allele can only occur in *Tbx1*-expressing cells and their descendants. The difference is likely to be due to incomplete Cre-induced recombination of the *Vegfr3*^*flox*^ allele in the critical cell population, which leaves sufficient VEGFR3 to support normal development in *Tbx1*^*Cre/+*^*;Vegfr3*^*flox/+*^ embryos. This interpretation is supported by the finding that the *Tbx1*^*Cre/+*^*;Vegfr3*^*flox/flox*^ genotype is associated with severe disruption of OFT development, indicating that the *Tbx1* expression domain is important for regulating *Vegfr3* in OFT development.

Where might the TBX1-*Vegfr3* interaction occur? The RNAscope experiments showed that *Tbx1* was activated in a modest number of differentiated ECs (*Vegfr3*-expressing) in the AS between E8.5 and E9.5. In contrast, when we traced *Tbx1* expressing cells and their descendents over time (48 hr, E8.5-E9.5), we observed extensive co-localization of the two proteins (GFP of the *Cre*-reporter and VEGFR3) in the AS at E9.5. Together, these results suggest that *Tbx1* is activated in EC progenitors, consistent with previous *in-vivo* and scRNA seq studies (Xu et al., 2005; Nomaru et al., 2021), perhaps in the CPM, the descendants of which populate the AS, and that It is also actively expressed in the EC after E8.5. As we did not observe expression of *Vegfr3* RNA or protein in the SHF, we propose that the genetic interaction affecting OFT development occurs in differentiated ECs in the AS around E9.5.

In conclusion, this study provides evidence of a regulatory mechanism involving *Tbx1* and *Vegfr3* that is critical for OFT morphogenesis. The results of the study point to a two-hit mechanism, whereby reduced dosage of *Vegfr3* in the entire expression domain (not only in the *Tbx1-Vegfr3* overlap domain) has a higher impact on OFT development when combined with *Tbx1* heterozygosity. The interaction between the two genes may be in part direct, in the region of expression overlap (aortic sac), and in part indirect, between the role of *Tbx1* in OFT morphogenesis, weakened by heterozygosity, and the endothelial-dependent role of *Vegfr3*, also weakened by heterozygosity.

At the molecular level, the vascular functions of *TBX1* have been attributed to interactions with molecules in the VEGF-C and Notch signaling pathways, (Chen et al., 2010; Cioffi et al., 2014; Martucciello et al., 2020). Based on previous studies demonstrating that NOTCH1 signaling in the endocardium is essential for outflow tract development (High et al., 2009), together with reports identifying dynamic VEGFR3-dependent endothelial populations during cardiovascular morphogenesis (Chen et al., 2014), we speculate that TBX1 may provide cues that are integrated by NOTCH1 in the endocardium, which in turn regulates endothelial programs including VEGFR3-mediated vascular remodeling during outflow tract morphogenesis.

## MATERIALS AND METHODS

### Mouse lines and genotyping

Mouse studies were performed in accordance with the animal protocol n°. 28/2022-PR, reviewed by the local IACUC committee and by the Italian Istituto Superiore di Sanità and approved by the Italian Ministero della Salute, according to Italian law and European guidelines. The following mouse lines were used: *Tbx1*^*Cre/+*^ (Huynh et al., 2007), *Vegfr3*^*flox/+*^ (Zarkada et al., 2015), *Vegfr3*^*+/−*^ (Martucciello et al., 2020), Rosa^*mTmG*^ (Muzumdar et al., 2007). Genotyping of mice was performed as in the original report.

### Immunofluorescence

Embryos were fixed in 4% paraformaldehyde, embedded in paraffin, and sectioned at thickness of 10µm. The sections were rehydrated, and antigen retrieval was performed in 10 mM sodium citrate pH6.0. Blocking was carried out using 0.5% milk,10% fetal bovine serum, and 1% bovine serum albumin H_2_O (blocking solution). Incubation with the primary antibody in blocking solution was carried out overnight at room temperature, and signal detection was performed using secondary antibodies conjugated with different fluorochromes (1:500). We used the following primary antibodies: anti-GFP (ab13970, Abcam; 1:1000); anti-Vegfr3 (14-5988-82, ebioscence; 1:400). The secondary antibodies were: Goat anti-Chicken IgY Alexa Fluor™ 488 Invitrogen A11039 (1:500) and Goat anti-Rat IgG (H+L) Alexa Fluor™ 594 Invitrogen A11007 (1:500). Imaging was carried out using a Leica Thunder Imaging System (Leica Microsystems) equipped with a Leica DFC9000GTC camera lumencor fluorescence LED lights source and the software Leica Application Suite X3.7.4.23463.

### RNA scope *in situ* hybridization

Whole embryos were fixed in 4% paraformaldehyde, dehydrated with increasing concentrations of methanol, and stored at 20°C. Embryos were hybridized with mRNA probes following the manufacturer’s instructions (RNAscope™, ACD-Biotechne); briefly, whole embryos were rehydrated in decreasing concentrations of methanol and permeabilized for 20 min with Protease III, followed by overnight incubation at 40°C with the probe solution (see list below). The signal was developed with TSA FITC (Akoya Bioscience, NEL741001KT; 1:500) and TSA-CY3 (Akoya Bioscience, NEL745001KT; 1:2000). The embryos were cryoprotected with increasing concentration of sucrose (10%, 20% and 30% sucrose in PBS 1×) at 4°C, OCT/sucrose 30% (50:50), embedded in OCT, cryosectioned (sagittal and transverse) at a thickness of 10µm and imaged using the Leica Thunder Imaging System described above. We used the following commercial probes provided by RNAscope™: Probe-Mm-Tbx1-C2 (481911-C2); Probe-Mm-VEGFR3 (481371). Imaging was carried out as for immunofluorescence experiments.

## Acknowledgments

We are grateful for the invaluable support provided by the Integrated Microscopy Core and the Animal Facility at the Institute of Genetic and Biophysics “ABT”/CNR, Naples. We thank Giuseppina Divisato at the Microscopy Core of the Department of Molecular Medicine and Medical Biotechnology, University of Naples Federico II; Lucia Mele for technical help with mouse handling and the microscopy core of the IGB-CNR. *Vegfr3*^*flox/+*^ mice were generously provided by Dr Kari Alitalo.

## Author Contributions

S Martucciello: data curation, formal analysis, validation, investigation, visualization, methodology and writing original draft. M Bilio: data curation, formal analysis, investigation, and methodology. S Cioffi: data curation, formal analysis, validation, investigation and methodology. M Cavallaro: data curation, formal analysis, investigation, methodology. A Baldini: conceptualization, supervision, funding acquisition. E Illingworth: conceptualization, supervision, funding acquisition, and writing original draft.

